# Bystro: rapid online variant annotation and natural-language filtering at whole-genome scale

**DOI:** 10.1101/146514

**Authors:** Alex V. Kotlar, Cristina E. Trevino, Michael E. Zwick, David J. Cutler, Thomas S. Wingo

**Author notes:** Corresponding author: Thomas S. Wingo, 505K Whitehead Building, 615 Michael Street NE, Atlanta, GA 30322-1047.

## Abstract

Accurately selecting relevant alleles in large sequencing experiments remains technically challenging. Bystro (*https://bystro.io/*) is the first online, cloud-based application that makes variant annotation and filtering accessible to all researchers for terabyte-sized whole-genome experiments containing thousands of samples. Its key innovation is a general-purpose, natural-language search engine that enables users to identify and export alleles and samples of interest in milliseconds. The search engine dramatically simplifies complex filtering tasks that previously required programming experience or specialty command-line programs. Critically, Bystro’s annotation and filtering capabilities are orders of magnitude faster than previous solutions, saving weeks of processing time for large experiments.

## Background

While genome-wide association studies (GWAS) and whole-exome sequencing (WES) remain important components of human disease research, the future lies in whole-genome sequencing (WGS), as it inarguably provides more complete data. The central challenge posed by WGS is one of scale. Genetic disease studies require thousands of samples to obtain adequate power, and the resulting WGS datasets are hundreds of gigabytes in size and contain tens of millions of variants. Manipulating data at this scale is difficult. To find the alleles that contribute to traits of interest, two steps must occur. First, the variants identified in a sequencing experiment need to be described in a process called annotation, and second, the relevant alleles need to be selected based on those descriptions in a procedure called variant filtering.

Annotating and filtering large numbers of variant alleles requires specialty software. Existing annotators, such as ANNOVAR[1], SeqAnt[2], VEP[3], and GEMINI[4] have played an important research role, and are sufficient for small to medium experiments (e.g.,10s to 100s of WES samples). However, they require significant computer science training to use in offline, distributed computing environments, and have substantial restrictions in terms of performance and the maximum size of the data they will annotate online. Existing variant filtering solutions are even more limited, with most analyses requiring researchers to program custom scripts, which can result in errors that impact reproducibility[5]. Therefore, annotation and filtering are not readily accessible to most scientists, and even bioinformaticians face challenges of performance, cost and complexity.

Here we introduce an application called Bystro that significantly simplifies variant annotation and filtering, while also improving performance by orders of magnitude and saving weeks of processing time on large data sets. It is the first program capable of handling sequencing experiments on the scale of thousands of whole-genome samples and tens of millions of variants online in a web browser, and integrates the first, to our knowledge, publicly-available, online natural-language search engine for filtering variants and samples from these experiments. The search engine enables real-time (sub-second), nuanced variant filtering, both across all samples and per sample, using simple phrases and interactive, web-based filters. Bystro makes it possible to efficiently find alleles of interest in any sequencing experiment without computer science training, improving reproducibility while reducing annotation and filtering costs.

## Results

To compare Bystro’s capabilities with other recent programs, we submitted 1000 Genomes[6] Phase 1 and Phase 3 VCF files for annotation and filtering (Figure 1). Phase 1 contains 39.4 million variants from 1,092 WGS samples, while Phase 3 includes 84.9 million alleles from 2,504 WGS samples. We first evaluated the online capabilities of the web-based versions of Bystro, wANNOVAR[7], VEP, and GEMINI (running on the Galaxy[8] platform). Bystro was the only program able to complete either 1000 Genomes Phase 1 or Phase 3 online, and was also the only application to handle a 6x10^6^ variant subset of Phase 3, a size representative of modest whole-genome experiments. When tested with 5x10^4^ – 1x10^6^ variant subsets of 1000 Genomes Phase 3, Bystro was approximately 144 – 212x faster than GEMINI/Galaxy in generating a downloadable annotation and searchable result database, and was significantly easier to use, as it did not require a separate annotation step (Figure 2). When tested on a small trio data set, Bystro was able to identify *de novo* variants without any additional software, and was 45x faster than GEMINI’s de_novo tool (Additional file 1: Table S1). Bystro and GEMINI/Galaxy produced similarly detailed outputs, with Bystro offering fewer, but more complete and recent sources, as well as more detailed annotations for some classes of data (Additional file 1: Table S2; Additional file 2). Notably GEMINI was found to work only with the hg19 human genome assembly, whereas Bystro supports hg19, hg38, and a variety of model organisms.

**Figure 1.**
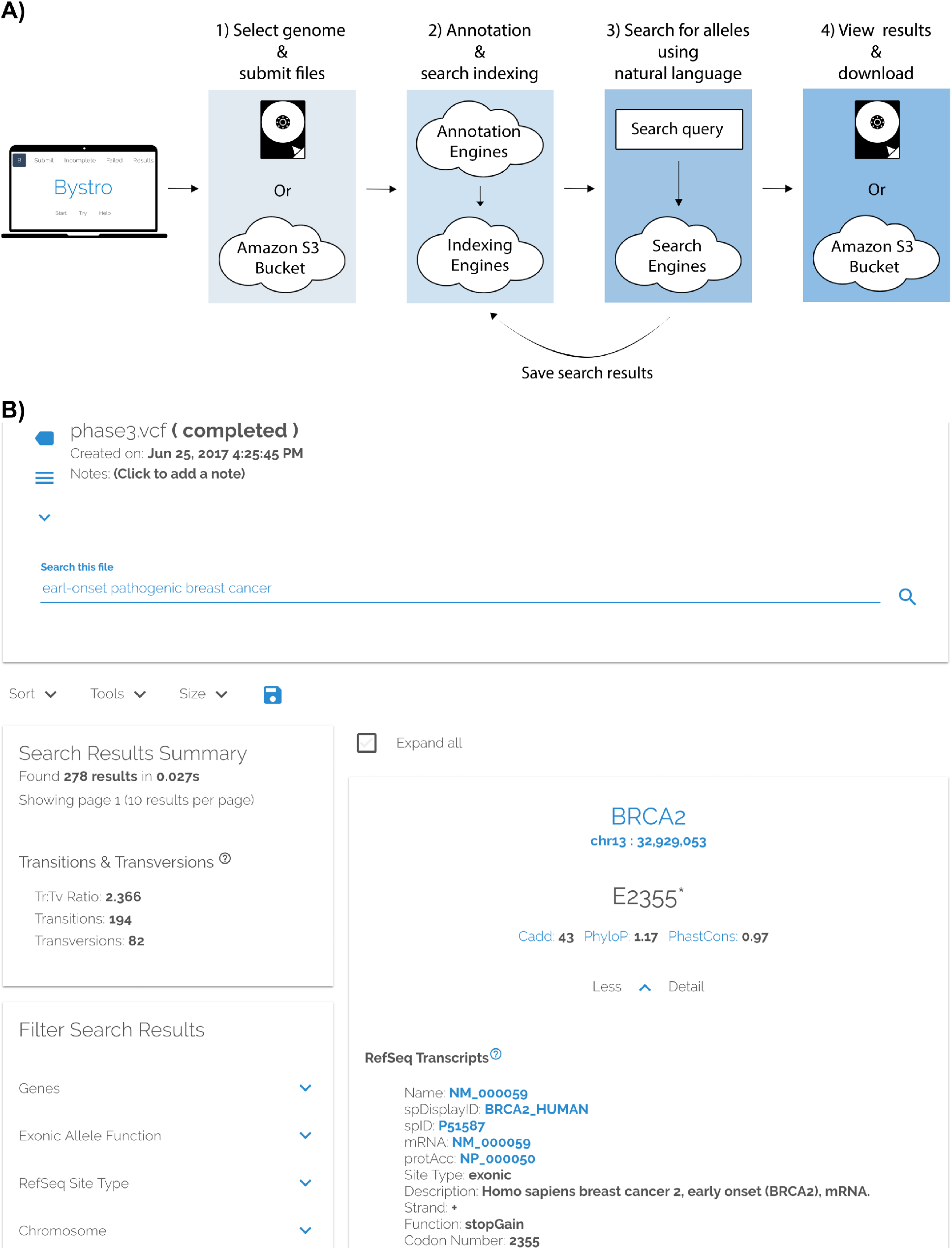
Using Bystro online to find alleles of interest in sequencing experiments. **A**) After logging in (*https://bystro.io/*), users upload one of more VCF or SNP-format files - containing alleles from a sequencing experiment - from a computer or a connected Amazon S3 bucket. Datasets of over 890GB, containing thousands of samples and tens of millions of variants are supported. The data is rapidly annotated in the cloud, using descriptions from public sources (e.g. RefSeq, dbSNP, Clinvar, and others). The annotated results can be filtered using Bystro’s natural-language search engine, and any search results can be saved as new annotations. Annotated experiments and saved results can be viewed online, downloaded as tab-delimited text, or uploaded back to linked Amazon S3 buckets. **B**) An example of using Bystro’s natural-language search engine to filter 1000 Genomes Phase 3 (*https://bystro.io/public*). To do so, users may type natural phrases, specific terms, numerical ranges, or apply filters on any annotated field. Queries are flexible, allowing misspelled terms such as “earl-onset” to accurately match. Complex tasks, such as identifying *de novo* variants can be achieved by using Boolean operators (AND, OR, NOT, +, -), exact-match filters, and user-defined terms. For instance, after labeling the “proband” and their “parents”, the user could simply search *proband –parents*, or combine with additional parameters for more refined queries, i.e. *proband –parents missingness* < .1 *gnomad.exomes.af_nfe* < *.001*.

We next tested offline performance on identical servers to gauge performance in the absence of web-related file-size and networking limitations. Bystro was 113x faster than ANNOVAR and up to 790x faster than VEP, annotating all 8.5x10^7^ variants and 2,504 samples from Phase 3 in less than 3 hours (Table 1). Furthermore, ANNOVAR was unable to finish either Phase 1 or Phase 3 annotations due to memory requirements (exceeding 60GB of RAM), and VEP annotated Phase 3 at a rate of 10 variants per second, indicating that it would need at least 98 days to complete. Critically, Bystro’s run time grew linearly with the number of submitted genotypes, suggesting that it could handle even hundreds of thousands of samples within days.

**Table 1.** Bystro, VEP, ANNOVAR offline command-line performance.

| Software | Dataset | Samples | Variants | Variants/s | Bystro vs |
| --- | --- | --- | --- | --- | --- |
| Bystro | 1000G Phase 3 chr1 | 2504 | $1 \times 10^6$ | $8156 \pm 195$ | - |
| | 1000G Phase 3 chr1 | 2504 | $2 \times 10^6$ | $8484 \pm 67.9$ | - |
| | 1000G Phase 3 chr1 | 2504 | $4 \times 10^6$ | $8516 \pm 57.2$ | - |
| | 1000G Phase 3 chr1 | 2504 | $6.5 \times 10^6$ | $7779 \pm 21.8$ | - |
| | 1000G Phase 1 | 1092 | $3.9 \times 10^7$ | $5417 \pm 76.8$ | - |
| | 1000G Phase 3 | 2504 | $8.5 \times 10^7$ | $7904 \pm 15.9$ | - |
| VEP | 1000G Phase 1 | 1092 | $3.9 \times 10^7$ | $18.67 \pm 0.58$ | 290x |
| | 1000G Phase 3 | 2504 | $8.5 \times 10^7$ | $10.00 \pm 0.00$ | 790x |
| ANNOVAR | 1000G Phase 3 chr1 | 2504 | $1 \times 10^6$ | $74.67 \pm 0.21$ | 109x |
| | 1000G Phase 3 chr1 | 2504 | $2 \times 10^6$ | $75.32 \pm 0.06$ | 113x |
| | 1000G Phase 3 chr1 | 2504 | $4 \times 10^6$ | $75.15 \pm 0.39$ | 113x |
| | 1000G Phase 3 chr1 | 2504 | $6.5 \times 10^6$ | NA | NA |
| | 1000G Phase 1 | 1092 | $3.9 \times 10^7$ | NA | NA |
| | 1000G Phase 3 | 2504 | $8.5 \times 10^7$ | NA | NA |

Bystro, VEP, and ANNOVAR were similarly configured with 8 threads on Amazon i3.2xlarge servers. “Dataset” refers to the VCF file used. “Variants/s” is the average of three trials. VEP performance was recorded after $2 \times 10^5$ sites in consideration of time. In runs of $1 \times 10^6$ or more annotated sites, VEP performance did not deviate from the $2 \times 10^5$ value. ANNOVAR could not complete the full Phase 1, Phase 3, or Phase 3 chromosome 1 datasets due to memory limitations. Thus, ANNOVAR was compared to Bystro on subsets of 1000 Genomes Phase 3 chromosome 1. Bystro run times included time taken to compress outputs. 1000 Genomes Phase 1 performance reflects IO limitations.

While offering significantly faster performance, Bystro also provided 3.5x the number of annotation output fields as ANNOVAR and 5.6x that of VEP (Additional file 3). Notably, unlike ANNOVAR or VEP, Bystro annotated each sample relative to its genotype, reporting homozygosity, heterozygosity, missingness, sample minor allele frequency, and labeling each sample as homozygous, heterozygous, or missing. In contrast, ANNOVAR provided only sample minor allele frequency, while VEP reported no sample-level data. We note that VEP is capable of providing per-sample annotations (heterozygosity/homozygosity status), but we were unable to use this feature for performance reasons. A detailed comparison of the exact settings used is given (Additional file 2; Additional file 3).

To investigate annotation accuracy, we next compared Bystro with ANNOVAR and VEP on a previously-analyzed synthetic dataset[9]. Overall, excellent concordance between all methods was noted (Additional files 4, 5, and 6). For instance, in comparison with ANNOVAR, allele position (>98%), allele identity (100%), and variant effects (>99%) were highly consistent across all classes of variation, for sites that Bystro did not exclude for quality reasons (Additional file 4).

In cases where the annotators disagreed, Bystro gave the more correct interpretations. For instance, Bystro and VEP excluded reference sites (ALT: “.”), while ANNOVAR annotated such loci as “synonymous SNV”; it is of course incorrect to call reference sites variant (Additional file 4; Additional file 5). In cases of insertions and deletions, which are often ambgiuously represented in VCF files due to the format’s padding requirements, Bystro always provided the parsimonious left-shifted representation, while ANNOVAR and VEP occasionally right-shifted variants (Additional file 4; Additional file 5). This is evident at chr15:42680000CA>CAA, where both ANNOVAR and VEP called the insertion as occuring after the first “A”, with 2 bases of padding, rather than the simpler option after the first base, “C”, with 1 base of padding (Additional file 1: Table S3). Similar results were found at multiallelic loci with complex indels (Additional file 1: Table S4).

Similarly, in cases where Bystro and ANNOVAR or VEP disagreed on variant consequences, Bystro always appeared correct relative to the underlying transcript set. For example, in the case of the simple insertion chr19:41123094G>GG, Bystro correctly identified all three overlapping transcripts (NM_003573;NM_001042544;NM_001042545), and noted the variant as coding (exonic) relative to all three. In contrast, ANNOVAR called the allele as disrupting a splice site, despite the fact that the nerest intron, and therefore splice site, was 37bp downstream (Additional file 1: Figure S1).

Additionally, Bystro’s strict VCF quality control measures substantially improved annotation accuracy. This is evident in the case of gnomAD, a VCF-format dataset that represents the largest experiment on human genetic variation. While Bystro and ANNOVAR provided identical gnomAD data for 93.7% of tested alleles, the remaining 6.3% were low-quality gnomAD results that were included in ANNOVAR and excluded from Bystro (Additional file 4). For instance, in the case of chr16:2103394C>T, ANNOVAR reported rs760688660, which failed gnomAD’s random forest qc step. We note that a 6.3% false-positive rate is similar to the frequency of common variation, and significantly larger than the frequency of rare variants, making ANNOVAR’s gnomAD annotations a potentially unreliable source of data for both common and rare variant filtering.

Next, we explored the Bystro search engine’s ability to filter the 84.9 million annotated Phase 3 variants. Bystro’s search engine was unique in its natural-language capabilities, and no other tested online program could handle the full Phase 3 dataset, or subsets as large as 6x10^6^ variants (Figure 2). First, we used Bystro’s search engine to find all alleles in exonic regions by entering the term “exonic” (933,343 alleles, 0.030 ± .001 seconds, Table 2). The search engine calculated a transition to transversion ratio of 2.96 for the query, consistent with previously observed values in coding regions. To refine results to rare, predicted deleterious alleles, we queried “cadd > 20 maf < .001 pathogenic expert review missense” (65 alleles, 0.029 ± 0.025s, Table 2). This search query could be written using partial words (“pathogen”), possessive nouns (“expert’s”), different tenses (“reviews”), and synonyms (“nonsynonymous”) without changing the results.

**Figure 2.**
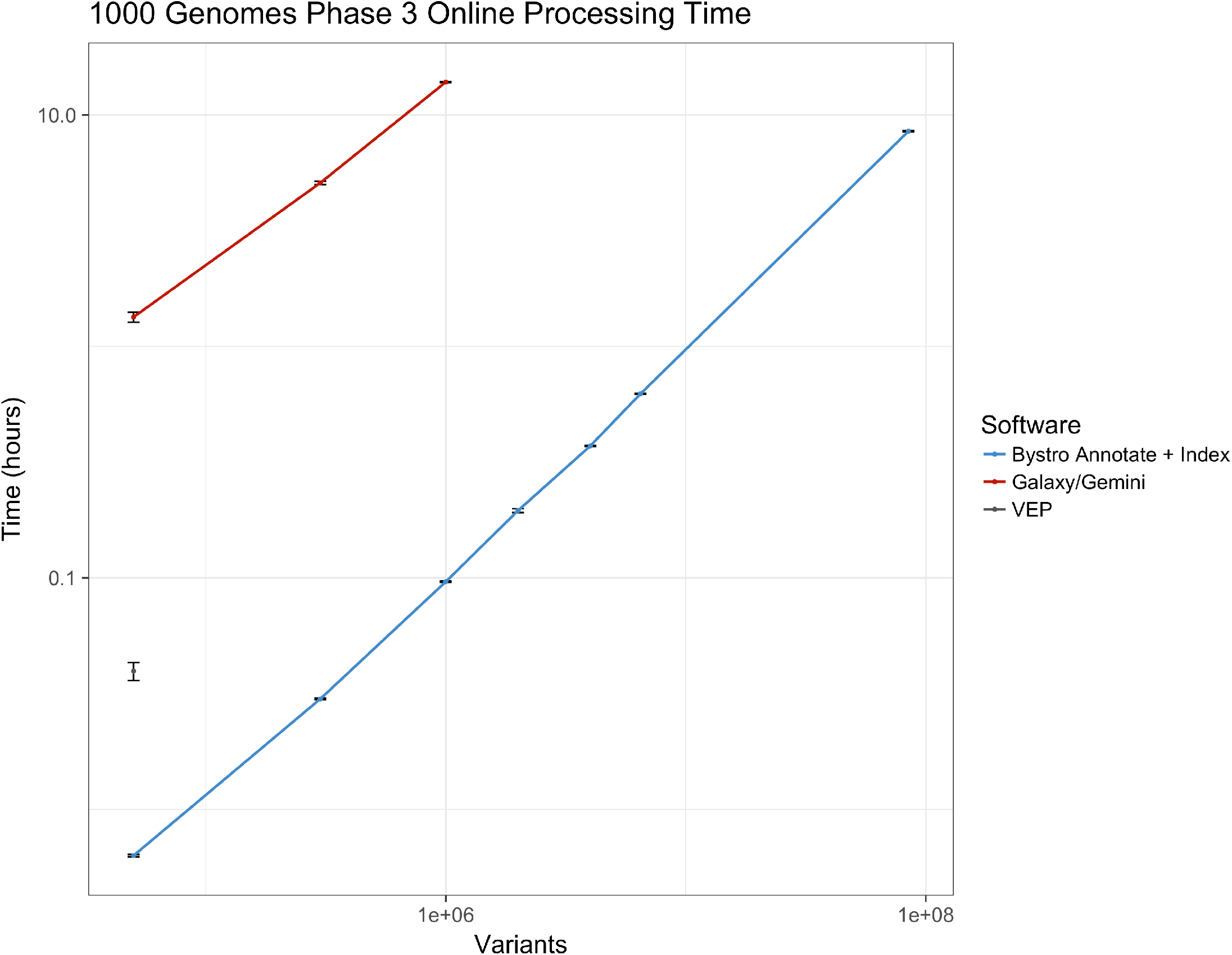
Online performance comparison of Bystro, VEP, wANNOVAR, and GEMINI. Bystro, wANNOVAR, VEP, and GEMINI (running on Galaxy) we run under similar conditions. Total processing time was recorded for 1000 Genomes Phase 3 WGS VCF files, containing either the full data set (2,504 samples, 8.49×10^7^ variant sites), or subsets (2,504 samples and 5×10^4^, 3×10^5^, 1×10^6^, and 6×10^6^ variants). Only Bystro successfully processed more than 1×10^6^ variants online: wANNOVAR (not shown) could not complete the smallest 5×10^4^ variant subset; VEP could not complete more than 5×10^4^ variants; and GEMINI/Galaxy could not complete more than 1×10^6^ variants. Online, VEP outputted a restricted subset of annotation data compared to its offline version. GEMINI and Bystro (but not VEP) outputted whole-genome CADD scores, while only Bystro also returned whole-genome PhyloP and PhastCons conservation scores. Bystro was faster than GEMINI/Galaxy by 144x-212x across all time points.

**Table 2.** Online comparison of Bystro and recent programs in filtering 8.49×10^7^ variants from 1000 Genomes.

| Group | Search query | Time (s) | Variants | Tr:Tv |
| --- | --- | --- | --- | --- |
| 1 | exonic | 0.030 ± 0.030 | 993,343 | 2.96 |
| 2 (a) | cadd > 20 maf < .001 pathogenic expert review missense | 0.029 ± 0.009 | 65 | 1.71 |
| 2 (b) | cadd > 20 maf < .001 pathogenic **expert’s review non-synonymous** | 0.036 ± 0.019 | 65 | 1.71 |
| 2 (c) | cadd > 20 maf < .001 **pathogen expert-reviewed nonsynonymous** | 0.044 ± 0.025 | 65 | 1.71 |
| 3 (a) | early onset breast cancer | 0.046 ± 0.029 | 4,335 | 2.51 |
| 3 (b) | **early-onset** breast cancer | 0.037 ± 0.020 | 4,335 | 2.51 |
| 3 (c) | **Early onset breast cancers** | 0.033 ± 0.015 | 4,335 | 2.51 |
| 4 (a) | Pathogenic nonsense Ehlers-Danlos | 0.038 ± 0.027 | 1 | NA |
| 4 (b) | **pathogenic** nonsense **E.D.S** | 0.078 ± 0.087 | 1 | NA |
| 4 (c) | **pathogenic stopgain eds** | 0.040 ± 0.022 | 1 | NA |

The full 1000 Genomes Phase 3 VCF file (853GB, $8.49 \times 10^7$ variants, 2,504 samples) was filtered in the publicly-available Bystro web application using the Bystro natural-language search engine. VEP, GEMINI, and wANNOVAR (not shown) were also tested, but were unable to annotate this data set or filter it. Bystro’s search engine uses a natural language parser that allows for unstructured queries: queries in groups 2, 3, and 4 show phrasing variations that did not affect results returned, as would be expected for a search engine that could handle normal language variation. “Tr:Tv” is the transition to transversion ratio automatically calculated for each query by the search engine. The transition to transversion ratio of 2.96 for the “exonic” query is close to the ~2.8-3.0 ratio expected in coding regions, suggesting that the search engine accurately identified exonic (coding) variants.

To test the search engine’s ability to accurately match variants from full-text disease queries, we first searched “early-onset breast cancer”, returning the expected alleles in *BRCA1* and *BRCA2* (4,335 variants, .037 ± .020s, Table 2). Notably, the queried phrase “early-onset breast cancer” did not exist within the annotation, and instead matched closely-related RefSeq transcript names, such as “Homo sapiens breast cancer 2, early onset (BRCA2), mRNA.” We next explored Bystro’s ability to handle synonyms and acronyms. To test the hypothesis that Bystro could interpret common ontologies, we queried “pathogenic nonsense E.D.S”, where “nonsense” is a common synonym for “stopGain” (a term annotated by the Bystro annotation engine), and “E.D.S” is an acronym for “Ehlers-Danlos Syndrome”. Bystro successfully parsed this query, returning a single *PLOD1* variant found in 1000 Genomes Phase 3 that introduces an early stop codon in all three of its overlapping transcripts, and which has been reported in Clinvar as “pathogenic” for “Ehlers-Danlos syndrome, type 4” (1 variant, .038s ± .027s, Table 2).

Since no other tested program could load or filter the 1000 Genomes Phase 3 VCF file online, we next compared Bystro to GEMINI (running on the Galaxy platform) on subsets of 1000 Genomes Phase 3. In contrast with GEMINI’s structured SQL queries, Bystro enabled shorter and more flexible searches. For instance, to return all missense, rare variants with CADD Phred scores larger than 15, GEMINI required a 162 character SQL query, while Bystro needed only 36 characters. Bystro also demonstrated synonym support, returning identical results for “missense” and “nonsynonymous” queries. Critically, Bystro’s search engine enabled real-time (sub-second) filtering, performing approximately four orders of magnitude faster than GEMINI on Galaxy while searching and returning similar volumes of data (Table 3).

**Table 3.**
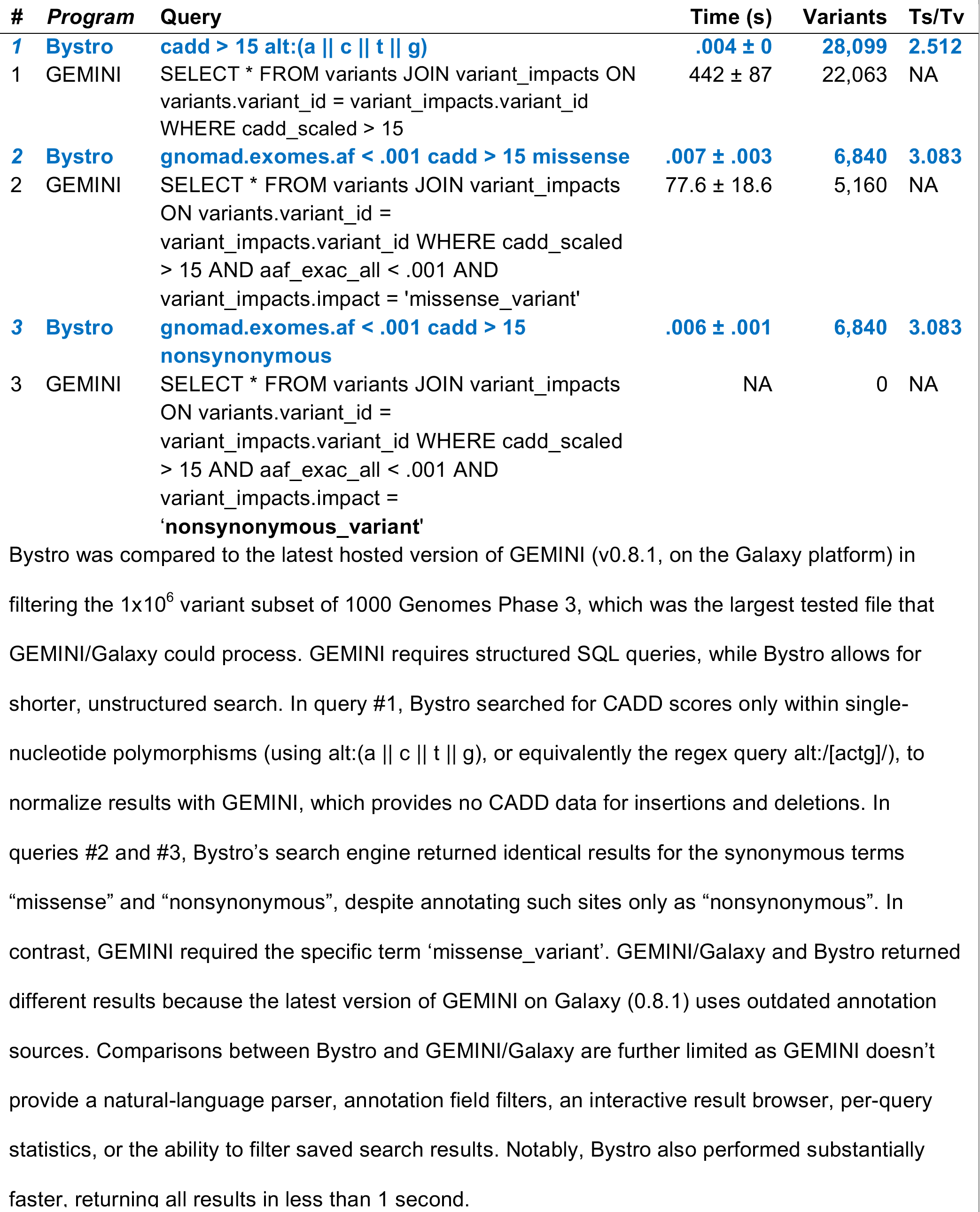
Online comparison of Bystro and GEMINI/Galaxy in filtering 1×10^6^ variants.

To test the accuracy of Bystro’s search engine relative to the underlying annotation, we first compared Bystro’s natural-language queries with Bystro’s “Filters”, which provide a complimentary, exact-match filtering option. All results were identical between the two methods (Additional file 1: Table S5). To control for the possibility that Bystro’s “Filters” were biased, we created separate Perl filtering scripts that searched for exact matches within the underlying tabdelimited text annotation. Again, results were completely concordant (Additional file 1: Table S5). Finally, to control for the possibility that both Bystro’s “Filters” and the Perl scripts were biased due to the programmer, we compared Bystro’s natural-language queries with Excel filters on a smaller dataset that could be manually examined. The queries were found completely specific in this comparison as well (Additional file 1: Table S6; Additional file 7).

## Discussion

The Bystro annotation and filtering capabilities are primarily exposed through a public web application (*https://bystro.io/*), and are also available for custom, offline installation. To ensure data safety, Bystro follows industry recommendations for password management, in-transit data security, and at-rest data security. Input and output files are encrypted at rest on Amazon EFS file systems, using AES 256-bit encryption, and every request for annotation or search data is authenticated by the web server using short-lived identity tokens. To further protect user data, annotation and search services are not directly open to the Internet, but require routing and authentication through the web server. Furthermore, all web traffic is encrypted using TLS (HTTPS), and password hashing follows the National Institute of Standards and Technology (NIST) recommended PBKDF2-HMAC-SHA512 strategy.

Creating an annotation online is as simple as selecting the genome and assembly used to make the variant call format (VCF)[10] or SNP[11] format files, and uploading these files from a computer or Amazon S3 bucket, which can be easily linked to the web application. Annotation occurs in the cloud, where distributed instances of the Bystro annotation engine process the data and send the results back to the web application for storage and display (Figure 1).

The Bystro annotation engine is open source, and supports diverse model organisms including *Homo sapiens* (hg19, hg38), *M. musculus* (mm9, mm10), *R. macaque* (rheMac8), *R. norvegicus* (rn6), *D. melanogaster* (dm6), *C. elegans* (ce11), S. *cerevisiae* (sacCer3). To annotate, it rapidly matches alleles from users’ submitted files to descriptions from RefSeq[12], dbSNP[13], PhyloP[14], PhastCons[14], Combined Annotation-Dependent Depletion (CADD), Clinvar[15], and gnomAD[16]. For custom installations, Bystro supports Ensembl, RefSeq, or UCSC Known Genes transcript sets, and can be flexibly configured include annotations from any files in genePredExt, wigFix, BED, or VCF formats.

The annotation engine is aware of alternate splicing, and annotates all variants relative to each alternate transcript. When provided sample information, Bystro also annotates all variants relative to all sample genotypes. In such cases, at every site it labels each sample as homozygous, heterozygous, or missing, and also calculates the heterozygosity, homozogosity, missingness, and sample minor allele frequency. Furthermore, in contrast with current programs that require substantial VCF file pre-processing, Bystro automatically removing low-quality sites, normalizes variant representations, splits multi-allelic variants, and checks the reference allele against the genome assembly. Critically, Bystro’s algorithm guarantees parsimonious (left-shifted) variant representations, even for multi-allelic sites containing complex insertions and deletions.

The Bystro annotation engine is designed to scale to any size experiment, offering the speed of distributed computing solutions such as Hail[17], but with less complexity. Current well-performing annotators - such as ANNOVAR and SeqAnt - load significant amounts of data into memory to improve performance. However, when these programs use multiple threads to take advantage of multicore CPUs they may exceed available memory (in some cases over 60GB), resulting in a sharp drop in performance or system crash. To solve this, Bystro annotates directly from an efficient memory-mapped database (LMDB), using only a few megabytes per thread, and because memory-mapped databases naturally lend themselves to the caching frequently accessed data, Bystro achieves most of the benefits of in-memory solutions, but without the per-thread penalties. This approach allows Bystro to take excellent advantage of multicore CPUs, while also enabling it to perform well on inexpensive, low-memory machines. Critically, when multiple files are submitted to it simultaneously, the Bystro annotation engine can automatically distribute the work throughout the cloud (or a user-configured computer cluster), gaining additional performance by processing the files on multiple computers (Figure 1). Furthermore, in reflection of the large sizes of both input sequencing experiments and the corresponding annotation outputs - on the order of terabytes for modern whole-genome experiments - Bystro accepts compressed input files, and directly writes compressed outputs. This ability to directly write compressed annotations with no uncompressed intermediate is critical given the rapid growth in sequencing experiment size.

When the web application receives a completed annotation, it saves the data and creates a permanent results page. Detailed information about the annotation, such as the database version used for the annotation is stored in a log file that the user may download. Users may then explore several quality control metrics, including the transition to transversion ratio on a per-sample or per-experiment basis. They may also download the results as tab-delimited text to their computer, or upload them to any connected Amazon S3 bucket. In parallel with the completion of an annotation, the Bystro search engine automatically begins indexing the results. Once finished, a search bar is revealed in the results page, allowing users to filter their variants using the search engine (Figure 1).

Unlike existing filtering solutions, Bystro’s Elasticsearch-based natural-language search engine accepts unstructured, “full-text” queries, and relies on a sophisticated language parser to match annotated variants. This allows it to offer the flexibility of modern search engines like Google and Bing, while remaining specific enough for the precise identification of alleles relevant to the research question. The Bystro search engine matches terms regardless of capitalization, punctuation, or word tense, and accurately finds partial terms within long annotation values. Like the annotation engine, the search engine is also exceptionally fast, automatically distributing indexed annotations throughout the cloud, enabling users to sift through millions of variants from large whole-genome sequencing experiments in milliseconds.

In order to provide flexible, but specific matches without relying on structured SQL queries, the search engine identifies the data type of every value in the annotation. Text undergoes stemming and lemmatization, which reduces the influence of grammatical variation, and is then tokenized into left-edge n-grams, which allows for flexible matching. Numerical data is stored in the smallest integer or float format that can accommodate it, allowing for rapid and accurate range queries. For complex queries, the search engine supports Boolean operators (AND, OR), regular expressions, and Levenshtein-edit distance fuzzy matches. It also has a built-in dictionary of synonyms, for instance equating “stopgain” and “nonsense”.

In some cases, text will match accurately, but not specifically; this most often happens with short, generic terms. For instance, querying “intergenic” alone may match the word “intergenic” in “long intergenic non-protein coding RNA” in refSeq’s description field, as well as “intergenic” in the refSeq’s siteType field. To help improve accuracy in such cases, Bystro provides three, closely related features: 1) “Aggregations” allows users to see the top 200 values for any text field, or equivalently the min, max, mean, standard deviation (and other similar statistics) for any numerical field. This allows users to quickly and precisely understand the composition of search results, as well as to generate summary statistics. 2) “Filters” allows users to refine queries, by forcing the inclusion or exclusion of any values found in any field. For instance, rather than query “intergenic”, it may be easier and more precise to simply click on the “refSeq.siteType” filter, and select the “intergenic” value. Any number of “Filters” may be combined with any natural-language query, containing up to 1 million words. 3) Bystro allows field names within a natural-language query for added specificity. For example, rather than searching for “intergenic”, the user could type “refSeq.siteType:intergenic”, to indicate that they wished to match “intergenic” specifically in the refSeq.siteType annotation field.

Bystro’s search engine also includes several features to increase flexibility beyond the contents of the annotation: 1) “Custom Synonyms” allows users to define their own terms and annotations. Among other uses, this make it is possible to label trios, which can be used to easily identify *de novo* variants and test allele transmission models. 2) “Search Tools” are small programs, accessible by a single mouse click, that dynamically modify any query to generate complex result summaries. Some of their functions include identifying compound heterozygotes. 3) “Statistical Filters” dynamically perform statistical tests on the variants returned from any query. For instance, the “HWE” filter allows users to exclude variants out of Hardy-Weinberg Equilibrium. This is an often-needed quality control step.

Most importantly, there is no limit to the number of query terms and “Filters” that can be combined, and users can save and download the results of any search query, which enables recursive filtering on a single dataset. The saved results are indexed for search, and hyperlinked to the annotations that they were generated from, forming permanent records that can be used to reproduce complex analyses. This multi-step filtering provides functionality similar to custom command-line filtering script pipelines, but is significantly faster, less error prone, and accessible to researchers without programming experience.

While Bystro’s annotation and filtering performance is currently unparalleled by any other approach, other software (such as Hail[17]) could achieve similar performance by implementing distributed computing algorithms like MapReduce[18], and spreading annotation workloads across many servers. Bystro demonstrates that these workarounds are unnecessary to achieve reasonable run-times for large datasets online or offline. Additionally, while Bystro’s natural-language search engine significantly reduces the difficulty of variant filtering, it does not handle language idiosyncrasies as robustly as more mature solutions like Google’s, and may return unexpected results when search queries are very short and non-specific, since such queries may have multiple correct matches. This is easily avoided by using longer phrases, by using “Custom Synonyms” to define more specific terms, by examining the composition of results using “Aggregations”, or by applying “Filters” to precisely filter results. Such considerations and options are well-documented in Bystro’s online user guide (*https://bystrio.io/help*).

## Conclusions

To date, identifying alleles of interest in sequencing experiments has been time-consuming and technically challenging, especially for whole-genome sequencing experiments. Bystro increases performance by orders of magnitude and improves ease of use through three key innovations: 1) a low-memory, high-performance, multithreaded variant annotator that automatically distributes work in cloud or clustered environments; 2) an online architecture that handles significantly larger sequencing experiments than previous solutions; and 3) the first publicly-available, general-purpose, natural-language search engine for variant filtering in individual research experiments. Bystro annotates large experiments in minutes, and its search engine is capable of matching variants within whole-genome datasets in milliseconds, enabling real-time data analysis. Bystro’s features enable practically any researcher – regardless of their computational experience - to analyze large sequencing experiments (e.g. thousands of whole-genome samples) within less than a day, and small ones (e.g. hundreds of whole-exome samples) in seconds. As genome sequencing continues the march toward ever-larger datasets and becomes more frequently used in diverse research settings, Bystro’s combination of performance and ease of use will prove invaluable for reproducible, rapid research.

## Methods

### Accessing Bystro

For most users, we recommend the Bystro web application (*https://bystro.io*), as it gives full functionality, supports arbitrarily large datasets, and provides a convenient interface to the natural-language search engine. Users with computational experience can download the Bystro open-source package (*https://github.com/akotlar/bystro*). Using the provided installation script or Amazon AMI image, Bystro can be easily deployed on an individual computer, computational cluster, or any Amazon Web Services (AWS) EC2 instance. Bystro has very low memory and CPU requirements, but benefits from fast SSD drives. As such we recommend at AWS instances with provisioned I/O EBS drives, RAID 0 non-provisioned EBS, or i2/i3-class EC2 instances.

Detailed documentation on Bystro’s use, as well as example search queries can be found at *https://bystro.io/help*.

## Bystro comparisons with ANNOVAR, wANNOVAR, VEP, and GEMINI/Galaxy

### Bystro Database

Bystro databases were created using the open-source package (*https://github.com/akotlar/bystro*). The hg19 and hg38 databases contains RefSeq, dbSNP, PhyloP, PhastCons, Combined Annotation-Dependent Depletion (CADD), and Clinvar fields, as well as custom annotations (Additional file 8). A complete listing of the original source data is enumerated in the Git repository (*https://github.com/akotlar/bystro/tree/master/config*). Other organism databases contain a subset of these sources, based on availability. Pre-built, up-to-date versions of these databases are publicly available (*https://github.com/akotlar/bystro*).

### WGS Datasets

Phase 1 and Phase 3 autosome and chromosome X VCF files were downloaded from *http://www.internationalgenome.org/data/*. Phase 1 files were concatenated using bcftools[19] “concat” function. Phase 3 files were concatenated using a custom Perl script (*https://github.com/wingolab-org/GenPro/blob/master/bin/mergeSnpFiles*). The Phase 1 VCF file was 895GB (139GB compressed), and the Phase 3 data was 853GB (15.6GB compressed). The larger size of Phase 1 can be attributed to the inclusion of extra genotype information (the genotype likelihood). The full Phase 3 chromosome 1 VCF file (6.4x10^6^ variants, 1.2GB compressed), and 5x10^4^-4x10^6^ variant allele subsets (8-655MB compressed) were also tested. All Phase 1 and Phase 3 data correspond to the GRCh37/hg19 human genome assembly. All data used are available (Additional file 9).

### Online annotation comparisons

For online comparisons, the latest online versions offered at time of writing were used. Bystro beta10 (September 2017), wANNOVAR (April 2017), VEP (April 2017), and GEMINI (Galaxy version 0.8.1, released February 2016, latest as of October 2017) were tested online with the full 1000 Genomes Phase 1 and Phase 3 VCF files, unless they were unable to upload the files due to file size restrictions (Additional file 2). Bystro was found to be the only program capable of uploading and processing the full Phase 1 and Phase 3 data sets, or subsets of Phase 3 larger than 1x10^6^ variants.

To conduct Bystro online annotations, a new user was registered within the public Bystro web application (*https://bystro.io/*). Phase 1 and Phase 3 files were submitted in triplicate, one replicate at a time, using the default database configuration (Additional file 2). Indexing was automatically performed by Bystro upon completion of each annotation. The Phase 3 annotation is publicly available to be tested (*https://bistro.io/public*).

The public Bystro server was configured on an Amazon i3.2xlarge EC2 instance. The server supported 8 simultaneous users. Throughout the duration of each experiment, multiple users had concurrent access to this server, increasing experiment variance, and limiting observed performance.

Online Variant Effect Predictor (VEP) submissions were done using the VEP web application (*http://www.ensembl.org/info/docs/tools/vep/index.html*). VEP has a 50MB (compressed) file size limit. Due to gateway timeout issues and this file size limit, data sets larger than 5x10^4^ variants failed to complete (Additional file 2).

Online ANNOVAR submissions were handled using the wANNOVAR web application. wANNOVAR could not accept the smallest tested file, the 5x10^4^ variant subset of Phase 3 chromosome 1 (8MB compressed) due to file size restrictions (Additional file 2).

Galaxy submission was made using the public Galaxy servers. Galaxy provides ANNOVAR, but its version of this software failed to complete any annotations, with the error “unknown option: vcfinput”. Annotations on Galaxy were therefore performed using GEMINI, which provides annotations similar to Bystro’s. Galaxy has a total storage allocation of 250GB (after requisite decompression), and both Phase 1 and Phase 3 exceed this size. Galaxy was therefore tested with the full 6.4x10^6^ variant Phase 3 chromosome 1 VCF file. Galaxy’s FTP server was able to upload the file; however, Galaxy was unable to load the data into GEMINI, terminating after running for 36 hours, with the message “This job was terminated because it ran longer than the maximum allowed job run time” (Additional file 2). Subsets of Phase 3 chromosome 1 containing 5x10^4^, 3x10^5^, and 1x10^6^ variants were therefore tested. Three repetitions of the 5x10^4^ variant submission were made. In consideration of the duration of execution, two repetitions were made of the 3x10^5^ and 1x10^6^ variants submissions. Since Galaxy does not record completion time, QuickTime was used to record each submission.

Bystro, VEP, and GEMINI online annotation times included the time to generate both a user-readable tab-delimited text annotation and a searchable database. GEMINI required an extra step to do so, using the query SELECT * FROM variants JOIN variant_impacts ON variants.name = variant_impacts.name.

### Variant filtering comparisons

After Bystro completed each annotation, it automatically indexed the results for search. The time taken to index this data was recorded. Once this was completed, the Bystro web application’s search bar was used to filter the annotated sequencing experiments. The query time, as well as the number of results and the transition to transversion ratio for each query, were automatically generated by the search engine and recorded. Query time did not take into account network latency between the search server and the web server. All queries were run six times and averaged. The public search engine, which processed all queries, was hosted on a single Amazon i3.2xlarge EC2 instance.

Since VEP, wANNOVAR, and Galaxy/GEMINI could not complete Phase 1 or Phase 3 annotations, variant filtering on these data sets could not be attempted. For small experiments VEP and GEMINI can filter based on exact matches, while wANNOVAR provides only pre-configured phenotype and disease model filters. VEP could annotate and filter at most only 5x10^4^ variants and was therefore excluded from query comparisons.

Galaxy/GEMINI was tested with subsets of 1000 Genomes Phase 3 of 1x10^6^ variants (the largest tested data set that Galaxy could handle), with the described settings (Additional file 2). In all GEMINI queries a JOIN operation on the variant_impacts table was used to return all variant consequences, and all affected transcripts, as Bystro does by default. Similarly, Bystro’s CADD query was restricted to single nucleotide polymorphisms (using alt:(A ║C║T║G)), as its behavior diverges from GEMINI’s at insertions and deletions: Bystro returns all possible CADD Phred scores at such sites, whereas GEMINI returns a missing value. Bystro returns all values to give users added flexibility: its search engine can accurately search within arrays (lists) of data. Furthermore, as GEMINI on Galaxy only provided the Ensembl transcript set, for all query comparisons with GEMINI, Bystro was configured to use Ensembl 90, which was the latest version available at time of revision. It is important to note that the latest version of GEMINI on Galaxy (0.8.1) dates to February 2016, and its databases are several years older: CADD (v1.0, 2014), Ensembl (v75, February 2014), ExAc (v0.3, October 2014), whereas Bystro uses up-to-date resources. As a result of searching more up to date Ensembl (v90), population allele frequency (gnomAD 2.0.1, the successor to ExAc 1.0), and CADD (v1.3) data, Bystro’s queries returned more data.

Since Galaxy does not report run times, QuickTime software was used to record each run, and the query time was calculated as the difference between the time the search submission entered the Galaxy queue, to the time that it was marked completed. Galaxy/GEMINI queries were each run more than 6 times. Because run times varied by more than 17x, the fastest consecutive 6 runs were averaged to minimize the influence of Galaxy server load.

All comparisons with the Bystro search engine are limited, because no other existing method provides natural-language parsing, and either rely on built-in scripts or require the user to learn a specific language (SQL).

### Filtering accuracy comparison

The latest version of Bystro (beta 10, September 2017) was used. For the 1000 Genomes query accuracy checks, the same underlying Ensembl-based Bystro annotation and search index was used as in the Bystro/GEMINI filtering comparison. Direct comparison to GEMINI were not made, in reflection of the age of the latest GEMINI Galaxy version (v0.8.1, with database sources dating to 2014). All Bystro queries from that comparison were saved, downloaded, and compared with Bystro “Filters”, which are exact-match alternatives to Bystro’s natural-language queries, as well as custom Perl filtering scripts that also require exact matches. A second query accuracy step was conducted, on the Yen et al 2017[9] VCF file. This file was annotated using the standard RefSeq Bystro database. The same queries used in the Bystro/GEMINI comparison were re-created on this smaller annotation, saved, downloaded, and compared with Bystro “Filters” and Excel filters. Excel filters were created in Excel 2016 (Mac), and required exact matches. All Excel-filtered and all Bystro query results were manually inspected for concordance (Additional file 7). All scripts generated and used in the comparison may be found at *https://github.com/akotlar/bystro-paper*.

### Offline annotation comparisons

To generate offline performance data, the latest versions of each program available at time of writing were used. Bystro beta10 (September 2017), VEP 86 (March 2017), and ANNOVAR (March 2017) were each run on separate, dedicated Amazon i3.2xlarge EC2 instances (Additional file 3). All programs’ databases were updated to the latest versions available as of March 2017 (VEP, ANNOVAR), or September 2017 (Bystro). All programs were configured to use the RefSeq transcript set.

Each instance contained 4 CPU cores (8 threads), 60GB RAM, and a 1920GB NVMe SSD. Each instance was identically configured. All programs were configured to as closely match Bystro’s output as possible, although Bystro output more total annotation fields (Additional file 3). Each data set tested was run 3 times. The annotation time for each run was recorded, and averaged to generate the mean variant per second (variant/s) performance. Submissions were recorded using the terminal recorder asciinema, and both memory and cpu usage were recorded using the **free** and **top** commands set to a 30 second timeout.

VEP was configured to use 8 threads and to run in “offline” mode to maximize performance, as recommended[3]. In each of three recorded trials, VEP was set to annotate from RefSeq and CADD, and to check the reference assembly (Additional file 3). Based on VEP’s observed performance, adding PhastCons annotations was not attempted. VEP’s performance was measured by reading the program’s log, which records variant/second performance every 5x10^3^ annotated sites. In consideration of time, VEP was stopped after at least 2x10^5^ variants were completed, and the 2x10^5^ variants performance was recorded.

ANNOVAR was configured to annotate RefSeq, CADD, PhastCons 100way, PhyloP 100way, Clinvar, avSNP, and ExAc version 0.3 (Additional file 3). ANNOVAR’s avSNP database was used in place of dbSNP, as recommended. We configured ANNOVAR to report allele frequencies from ExAc, because it does not do so from either avSNP or dbSNP databases. When annotating Phase 1, Phase 3, or Phase 3 chromosome 1, ANNOVAR crashed by exceeding the available 60GB of memory. It was therefore tested with the subsets of Phase 3 chromosome 1 that contained 1x10^6^ – 4x10^6^ variants.

Bystro was configured to annotate descriptions from RefSeq, dbSNP 147, CADD, PhastCons 100way, PhyloP 100way, Clinvar, and to check the reference for each submitted genomic position (Additional file 3).

### Annotation accuracy comparison

The latest version of Bystro (beta 10, September 2017), ANNOVAR (July 2017), and VEP (version 90) at the time of revision submission were used. All programs’ databases were updated to the latest version available. RefSeq-based databases were downloaded using each program’s database builder. All programs were compared on the Yen et al 2017 VCF file [9] for position, variant call, and variant effects, based on each programs’ respective RefSeq database. The Yen et al VCF file *fileformat* header line was modified to “VCFv4.1” to allow programs to recognize it as a valid VCF file. This modified file is available: https://github.com/akotlar/bystro-paper. For the SnpEff comparison, annotations were adapted from Additional File 1 of Yen et al 2017[9]. ANNOVAR was additionally configured with gnomAD genomes, gnomAD exomes, and CADD 1.3, and compared to Bystro on the corresponding values.

### Additional Files

Additional file 1: This file contains 1) a feature comparison of tested programs, 2) investigation of annotation concordance between tested programs, 3) investigation of Bystro query accuracy (.docx, 1.4MB)

Additional file 2: Description of online comparison settings (.xlsx, 859KB)

Additional file 3: Description of online comparison settings (.xlsx, 40KB)

Additional file 4: Bystro vs ANNOVAR annotation comparison details (.xslx, 87KB)

Additional file 5: Bystro vs VEP annotation comparison details (.xslx, 701KB)

Additional file 6: Bystro vs SnpEff annotation comparison details (.xslx, 63KB)

Additional file 7: Bystro queries vs Excel filters concordance details (.xslx, 166KB)

Additional file 8: Species supported at time of writing, and their configurations (.xslx, 36KB) Additional file 9: URLs of 1000 Genomes Phase 1, 1000 Genomes Phase 3, and Yen et al 2017 VCF files used (.xslx, 47KB)

## Declarations

### Availability of data and materials

The Bystro web application is freely accessible at *https://bystro.io/*, and features detailed interface documentation (*https://bystro.io/help*). The Bystro annotator, search indexer, distributed queue servers, and database builder source code is freely available on GitHub (*https://github.com/akotlar/bystro*) and Zenodo (doi: *10.5281/zenodo.1012417*), under the Apache 2 open-source license [20]. The software is written in Perl and Go programming languages and runs on Linux and Mac operating systems. Detailed documentation for Bystro software is provided at *https://github.com/akotlar/bystro/blob/master/README.md*. The datasets generated during and/or analyzed during the current study are available in the GitHub repository, *https://github.com/akotlar/bystro-paper* [6],[9].

### Author contributions

A.V.K designed, wrote, and tested Bystro and performed experiments. C.E.T wrote Bystro documentation and performed quality control. M.E.Z and D.J.C. contributed to the design of Bystro and experiments. T.S.W. designed and wrote Bystro and designed and performed experiments. A.V.K. and T.S.W. wrote the manuscript with contributions from all authors.

## Acknowledgements

We thank Kelly Shaw and Katherine Squires for beta testing and design suggestions. We thank Viren Patel and the Emory Integrated Genomics Core (EIGC) for technical support.

## Ethics approval and consent to participate

Not applicable

